# 3D (x-y-t) Raman imaging of tomato fruit cuticle: microchemistry during development

**DOI:** 10.1101/2022.06.01.494410

**Authors:** Ana González Moreno, Eva Domínguez, Konrad Mayer, Nannan Xiao, Peter Bock, Antonio Heredia, Notburga Gierlinger

## Abstract

The cuticle of tomato fruits was studied *in-situ* using Confocal Raman Microscopy. Microsections from cuticles isolated at different developmental stages were scanned to reveal the distribution of cuticle components with a spatial resolution of 342 nm by univariate and multivariate data analysis. From the three main components, cutin, polysaccharides and aromatics, the latter one exhibit the strongest Raman scattering intensity. Therefore, Raman imaging opened the view on phenolic acids and flavonoids within the cuticle and resulted in three schematic cuticle models depicting development.

At the earliest stage of development, which corresponded to the procuticle layer, phenolic acids were found across the entire cuticle. Based on a mixture analysis with reference component spectra, the phenolic acids were identified as mainly esterified *p*-coumaric acid together with free *p*-hydroxybenzoic acid. Later in development, during the cell expansion period of growth, phenolic acids accumulated in an outermost layer of the cuticle and in the middle region of the pegs. In these stages of development cellulose and pectin were appeared towards the epidermal layer, where later during ripening the flavonoid impregnation started. In the first ripening stage chalconaringenin was observed, while methoxylated chalcones were chosen by the algorithm to fit the mature cuticle spectra. The co-location with carbohydrates and esterified *p*-coumaric acid and methoxylated chalconaringenin suggest that they link polysaccharide and cutin domains. Within the cutin matrix, aromatics confer mechanical and thermal functions, while the outermost phenolic acid layer displays UV-B protection of the plant tissue.

**One-sentence summary:** New insights into the distribution of cutin, carbohydrates and phenolics along cross sections of green and mature tomato fruit cuticles by Raman mapping and multivariate data analysis.

## Introduction

The plant cuticle is the outermost barrier, which covers the surface of aerial non-lignified organs (leaves, stems, flowers and fruits) of land plants. This outer layer is responsible for some critical functions in the plant kingdom such as providing mechanical support and defence against pathogens, radiation or mechanical injuries, mediating water and gas exchange with the environment and preventing from organ adhesion during first stages of development (Martin and Rose, 2014). Tomato fruit cuticle is one of the most studied among species due to its economic importance, ample availability of genetic and genomic resources, together with the easy isolation of its cuticle, lack of stomata and feasibility to perform mechanical and permeability tests.

The main component of the tomato fruit cuticle is the amorphous polymeric matrix named cutin, whose main monomers are 9(10),16-dihydroxyhexadecanoic acids (Heredia, 2003). As also cell wall derived polysaccharides are present (López-Casado et al., 2007; Philippe et al., 2020), the cuticle has been defined as a cutinized cell wall (Domínguez et al., 2011). Additionally, waxes are present with around 3-5% in the tomato fruit cuticle (Domínguez et al., 2009a) and phenolics with around 0.8-7.5% depending on the stage of development (España et al., 2014b). Waxes can be deposited on the surface (epicuticular waxes) or embedded within the cuticle matrix (intracuticular waxes). The phenolic fraction identified in the tomato fruit cuticle is composed of cinnamic acid derivatives such as *p*-coumaric acid and *p*-hydroxybenzoic acid, and the flavonoid chalconaringenin (Lara et al., 2019). Despite their presence as minor components, several physiologically relevant functions have been ascribed to the wax and phenolic fractions of the cuticle, such as water permeability, thermal properties and mechanical performance (Schreiber and Schönherr, 1990; Casado and Heredia, 2001; Vogg et al., 2004; Khanal et al., 2013; España et al., 2014b; España et al., 2014a; Jetter and Riederer, 2016; Benítez et al., 2022). Additionally, the polysaccharide fraction has been shown to play an important mechanical role (López-Casado et al., 2007).

The chemical composition of the tomato fruit cuticle is known (Hunt and Baker, 1980; Vogg et al., 2004). But, there is a knowledge gap on the spatial distribution of each component due to the lack of spatial resolution of wet chemical methods, no stains and/or antibodies are available for specific components, and the fact that cutin matrix itself masks other components and impedes penetration (Fernández et al., 2016; Segado et al., 2016). Determining the location and distribution of each component is crucial to understand their influence on the overall biophysical and macroscopic properties of the cuticle. Flavonoids have been shown to form clusters within the cuticle (Domínguez et al., 2009b). They have been postulated to act as fillers providing mechanical resistance (España et al., 2014b; España et al., 2014a), similar to the role suggested for intracuticular waxes (Khanal et al., 2013). However, it remains undetermined whether these compounds are specifically located at a certain region of the cuticle or distributed throughout. Polysaccharides, on the other hand, have been traditionally assumed to be present in the inner side of the cuticle, whereas the upper section was understood as a cutin matrix devoid of polysaccharides (Fernández et al., 2016). In tomato, polysaccharides have been found to be an important fraction (25-30% cuticle) (Domínguez et al., 2008). However, immunolocation of polysaccharides within the tomato fruit cuticle has been unsuccessful (Segado et al., 2016). Only after a partial degradation of the cutin matrix, the polysaccharide domain has been detected in the cuticle of several species, showing a location across the entire cuticle width (Guzmán et al., 2014a; Guzmán et al., 2014b).

Confocal Raman Microscopy (CRM) enables to reveal the distribution of plant components on the micro-scale (Gierlinger and Schwanninger, 2006; Richter et al., 2011; Mansfield et al., 2013; Mateu et al., 2016; Bock et al., 2021; Sasani et al., 2021) since it allows an *in situ* chemical resolution of around 300 nm (Prats-Mateu et al., 2018a). It is a rapid and non-destructive technique with simple sample preparation. The location of several components is possible simultaneously, but the multicomponent nature of biological samples results in broad and overlapping bands. To assign every Raman band to a specific component may become difficult and previous knowledge of the chemical composition becomes helpful, as well as infrared spectroscopy studies. Infrared, as the complementary vibrational spectroscopy technique, has been widely used to study the composition and structural characteristics of the cuticle (Mazurek et al., 2013; Heredia-Guerrero et al., 2014; Farber et al., 2019b; Farber et al., 2019a) and hence, numerous bands have been assigned to different cuticle components. The intensity of some of these bands can be used to image the assigned specific components by univariate approaches (Gierlinger 2018). However, the band overlap in plant samples often needs to be overcome by multivariate approaches to retrieve a maximum of information on microchemical variation within the tissue (Gautam et al., 2015; Szymańska-Chargot et al., 2016).

In this work, cuticle components were tracked on the microscale within the tomato fruit cuticle using CRM. We revealed the chemical microscale heterogeneity by univariate and multivariate data analysis. To find the purest component spectra non-negative matrix factorization was used and to confirm specific components or reveal their mixture with other components an orthogonal matching pursuit algorithm in combination with a reference library was applied. With these methodologies we aim to 1) discriminate the main components, lipids, carbohydrates and phenolics as well as 2) identify specific molecules e.g. benzoic, cinnamic acids and flavonoids and 3) image their quantity and location across the cuticle and 4) throughout cuticle development.

## Results

### Band assignment to different cuticle components and their changes during development

Figure 1 shows average Raman spectra of cross sections of isolated cuticles from 8 to 55 days after anthesis (daa). Raman spectra of cuticles at early stages of development show more noise and lower intensity, followed by a general increase in band intensity during development. Additionally, a slight shift of the Raman bands towards lower wavenumbers was observed during ripening (Fig. 1).

**Figure 1.**
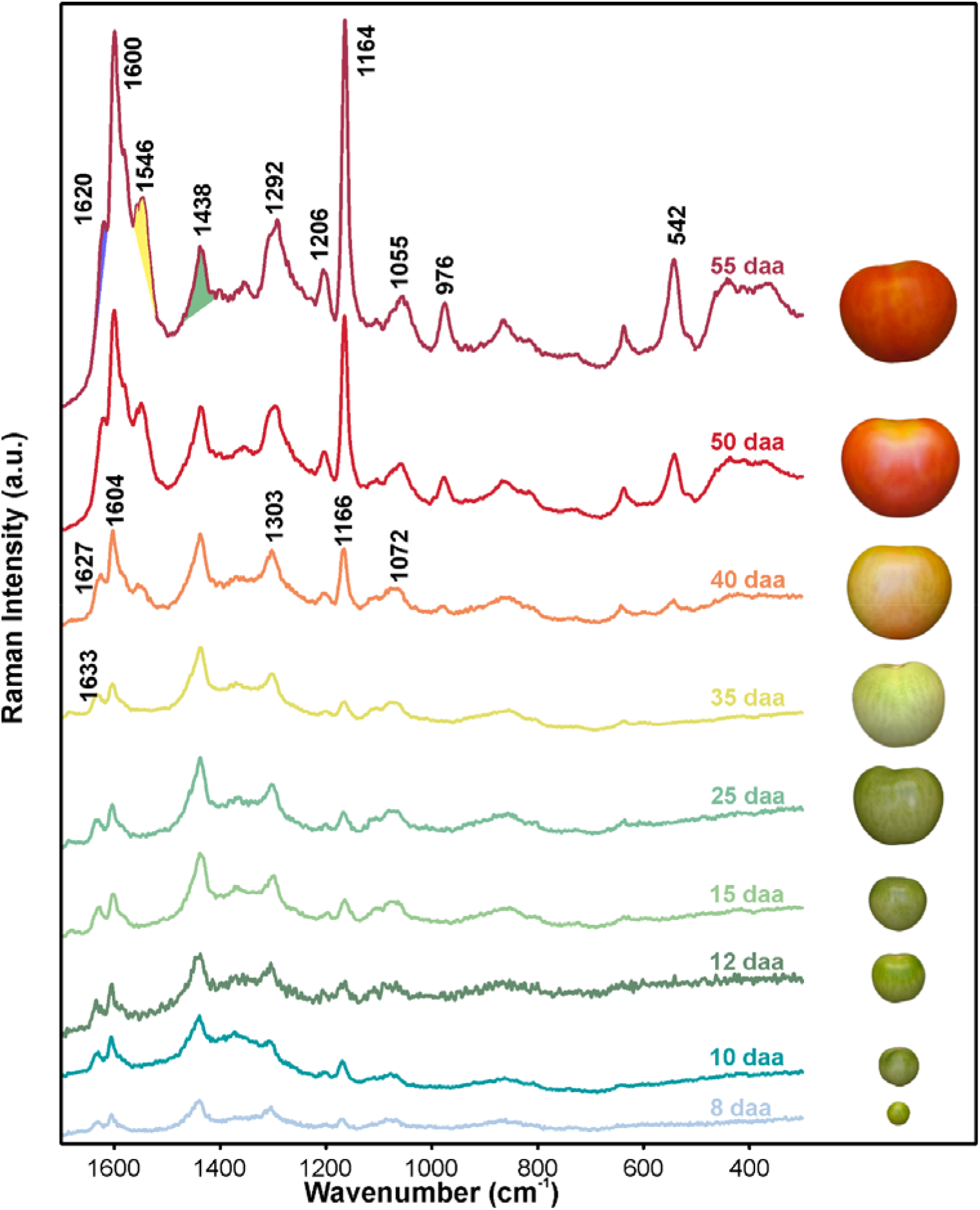
Average Raman spectra of tomato fruit cuticle at different stages of development. Spectral signatures of ‘Cascada’ fruit cuticles from 8 to 55 daa (days after anthesis) and their photographs on the right (taken from (Segado et al., 2016). Band positions of characteristic component bands are highlighted (in cm^−1^) and their assignments are listed in Table 1.

In the Raman spectra, the bands representative for specific components are highlighted (Fig. 1) and their band assignment is given in Table 1. The plant polyester cutin is reflected by the bands at 1438, 1278, 1104 and 1164 cm^−1^ (see Table 1 for assignments). Although they are representative of cutin, they can overlap with other components present in tomato fruit cuticle, mainly polysaccharides and waxes (Prinsloo et al., 2004; Heredia-Guerrero et al., 2014; Gierlinger, 2018; Farber et al., 2019a). Nevertheless, due to the higher Raman scattering intensity of polyester in comparison with signals from polysaccharides and the comparatively low amount of waxes in tomato fruit cuticle, mainly cutin contributes to these bands. Polysaccharides display strong bands at e.g. 1095, 1122 cm^−1^ (Cellulose, (Wiley and Atalla, 1987) and between 853-856 cm^−1^ (pectin, (Synytsya et al., 2003)), but these signals have low intensity compared with those from other cuticle components such as cutin or phenolics and are consequently partially masked. Bands at 1055, 1060, 1132, 1292 and 1438 cm^−1^ (Table 1) are characteristic of cuticle waxes, but their low amount (~3% of total weight) (Domínguez et al., 2008) hampers their selective identification.

**Table 1.**
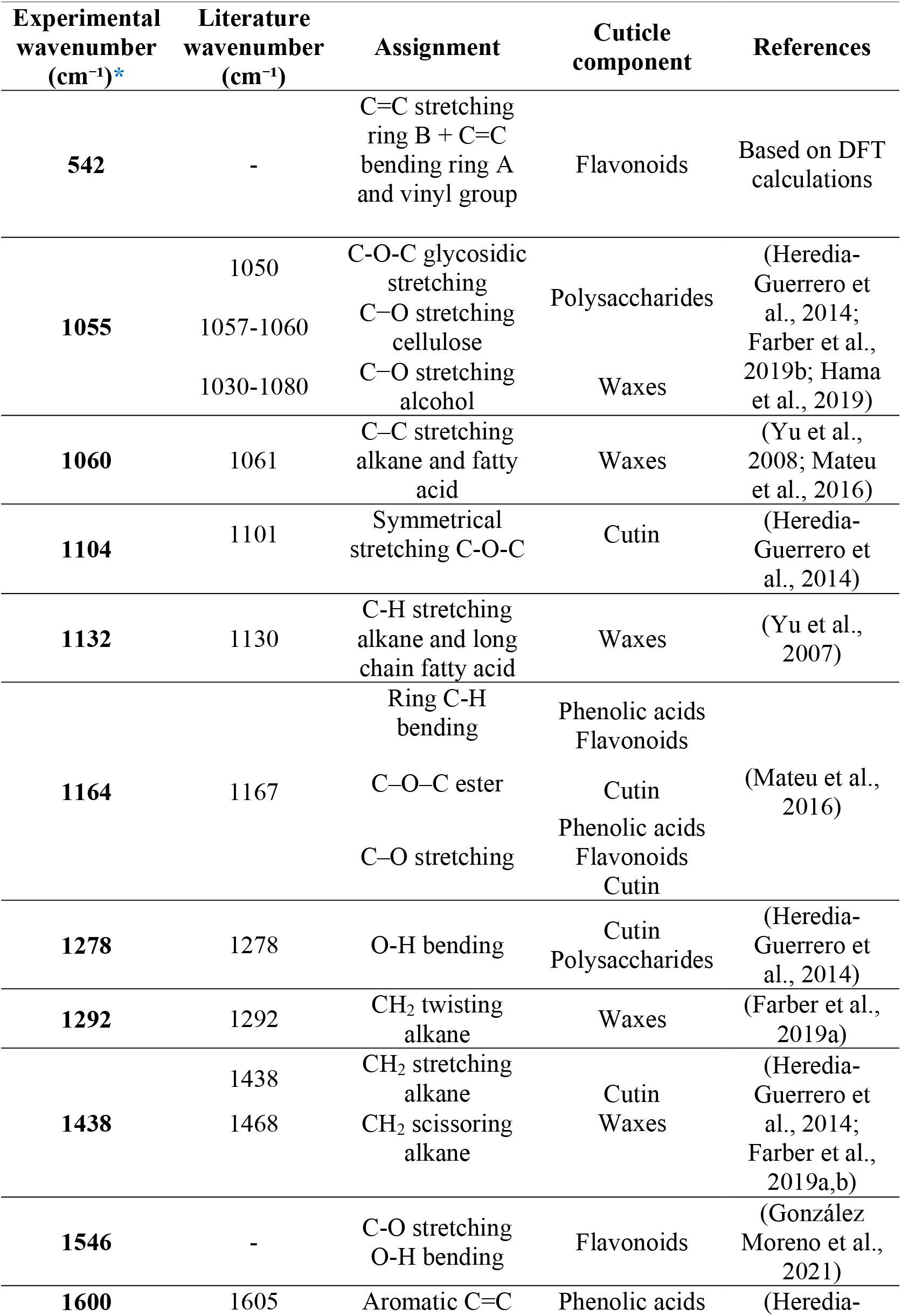
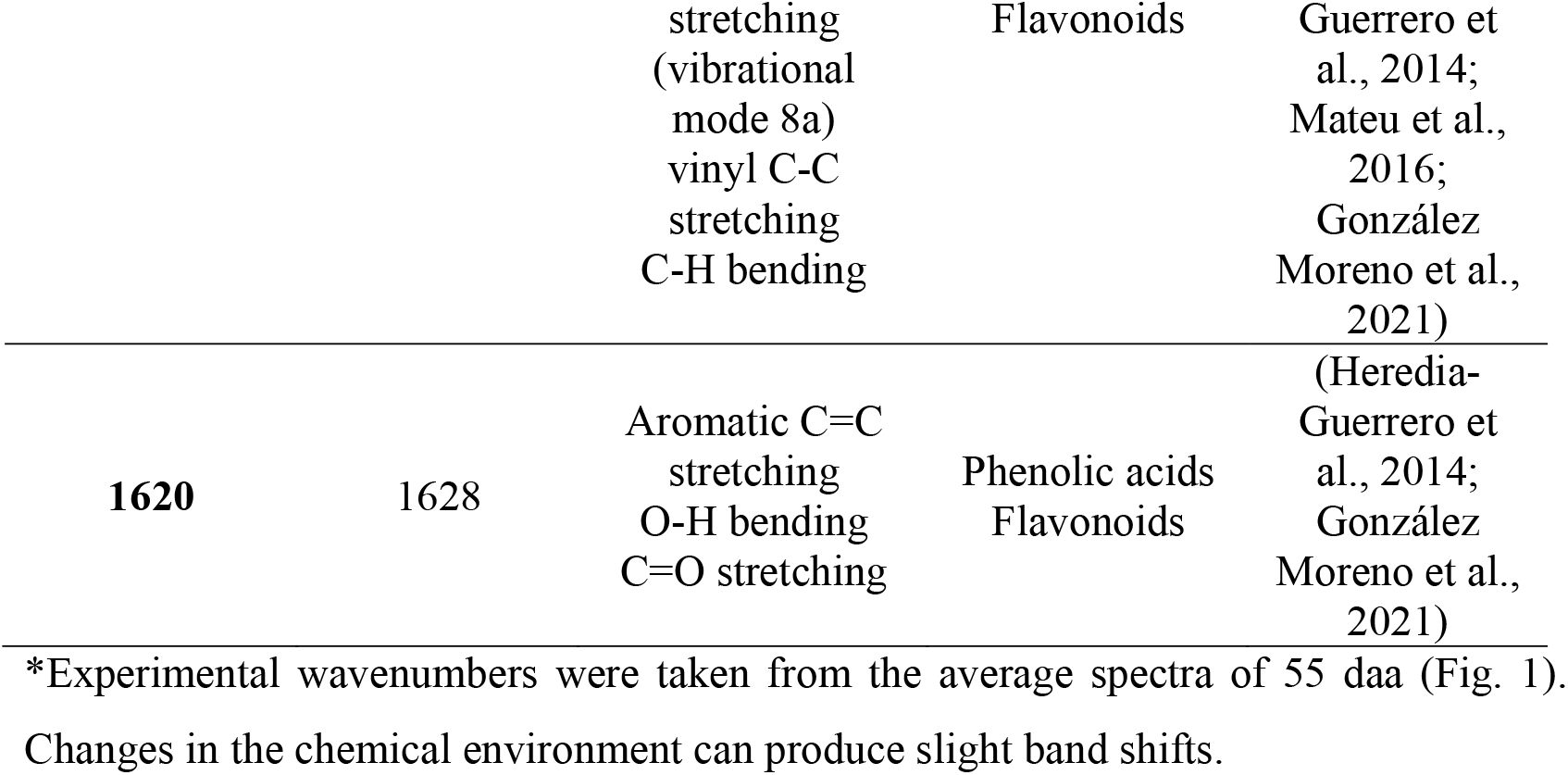
Raman band assignments according to literature and DFT calculations (see Table S1 for more details) show the multicomponent nature and problem of overlapping bands.

Phenolic compounds, on the other hand, exhibit strong signals in Raman spectroscopy. Tomato fruit cuticle accumulates two classes of phenolic compounds: hydroxycinnamic acid derivatives, mainly *p*-coumaric acid and *p*-hydroxybenzoic acid, present throughout development, and the flavonoid chalconaringenin that only accumulates during ripening and confers an orange-yellow colour to the cuticle (España et al., 2014b). Signals centred at around 1164 and 1600 cm^−1^ (Fig1, see Table 1 for assignment) are common for both types of phenolic compounds due to their structural similarities. The band centred around 1630 cm^−1^ was selected to monitor phenolic acid derivatives, as it is strong in coumaric acid spectra and much weaker in flavonoid (chalconaringenin) spectra (Fig.S1). On the other hand, bands centred around 542 and 1546 cm^−1^ appear only during ripening (Fig.1) and can be exclusively attributed to chalconaringenin (Fig. S1). *DFT* calculations of free chalconaringenin revealed the nature of their bands centred at 546, 1567 and 1509 cm^−1^ in its experimental spectrum (Fig.S1). The band centred around 546 cm^−1^ (542 cm^−1^ in tomato fruit Raman spectrum) is attributed to stretching of the ring B coupled to C=C bending of the ring A and the vinyl group (see Table S1). No band centred at 1546 cm^−1^ (the emerging band in the ripe tomato fruit cuticle) was detected for free naringenin chalcone; nevertheless, in this region two bands are exhibited by the flavonoid: one centred at 1567 cm^−1^ assigned to the stretching of the C=O coupled with the in-plane O-H bending of the H bonding (González Moreno et al., 2021); and another centred at 1509 cm^−1^ corresponded to the C=O stretching and O-H bending (H bond) coupled with other ring vibrations (see Table S1 for more details).

Plotting the band area of ~1630, 1546 and 1438 cm^−1^ revealed semi-quantitatively the progression of the phenolic acids, flavonoids and cutin enriched matrix, respectively (Fig. 2A). During the growing period, from 8 to 35 daa the similar intensity of the band at 1630 cm^−1^ suggested constant phenolic acid contents, while later during ripening an increase was observed. Also the flavonoid band 1546 cm^−1^ increased strongly during ripening from 40-55 daa, but was not present during the growing period. The 1438 cm^−1^ band area increased during the 8-15 daa period and then remained more or less constant until red ripe (55 daa). To level out changes due to different focal plane or overlapping band intensities, ratios between the band areas of phenolic acids and cutin as well as between flavonoids and phenolic acids were additionally calculated (Fig. 2B). The phenolic acids/cutin ratio remained quite constant during development (including a small increase during ripening) and suggested a co-occurrence of the two from the very beginning. The flavonoid/phenolic acid ratio, on the other hand, clearly indicated that flavonoids mainly contributed to the increase in phenolic compounds during ripening and that their influence on the phenolic acid band (1630 cm^−1^) is very small.

**Figure 2.**
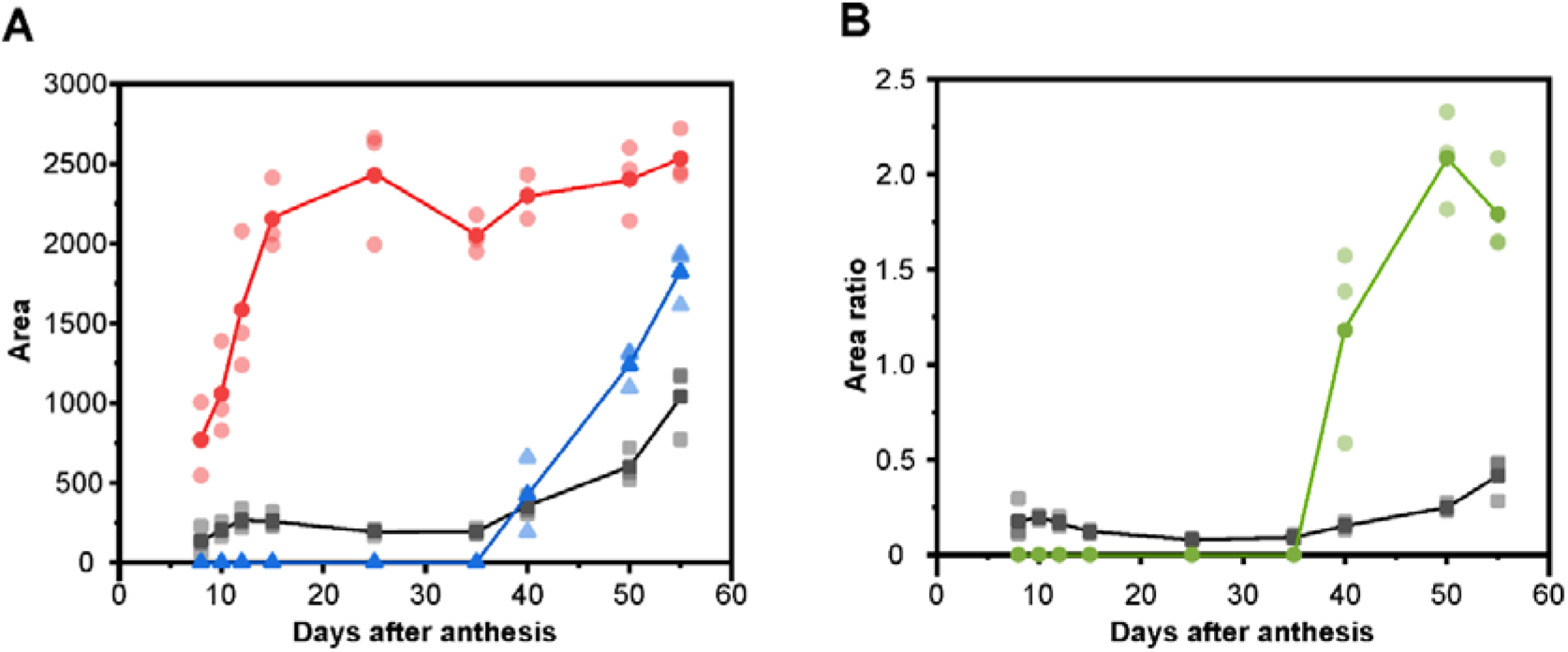
Changes in three selected Raman band areas during development. **(A)** Development of the 1630, 1546 and 1438 cm^−1^ Raman band areas (see Fig. 1) associated with the presence of phenolic acids (grey squares), flavonoids (blue triangles) and cutin enriched matrix (red circles) in isolated tomato ‘Cascada’ fruit cuticles from 8 to 55 daa. Dark symbols represent mean values and the pale ones in the same colour correspond to the different replicates. **(B)** Ratios of the phenolic acids/cutin (1630/1438; grey squares) and flavonoid/phenolic acid (1546/1630; green circles) band areas.

### Imaging the spatial distribution of tomato fruit cuticle components by band integration

To visualize the spatial distribution of the cuticle components within the cross section of the cuticle, the bands assigned to cutin, phenolic acids and flavonoid were integrated. The thickness of the 8 daa sample was below the limit of resolution of the technique and could not be studied with the imaging approach. Figure 3 shows the results of this univariate analysis in four selected stages of development. Raman images of the cutin enriched matrix depicted a homogenous distribution throughout the entire section of the cuticle in all stages of development, without any detectable differences in intensity between the periclinal section of the cuticle and the pegs (cutinized anticlinal epidermal wall).

**Figure 3.**
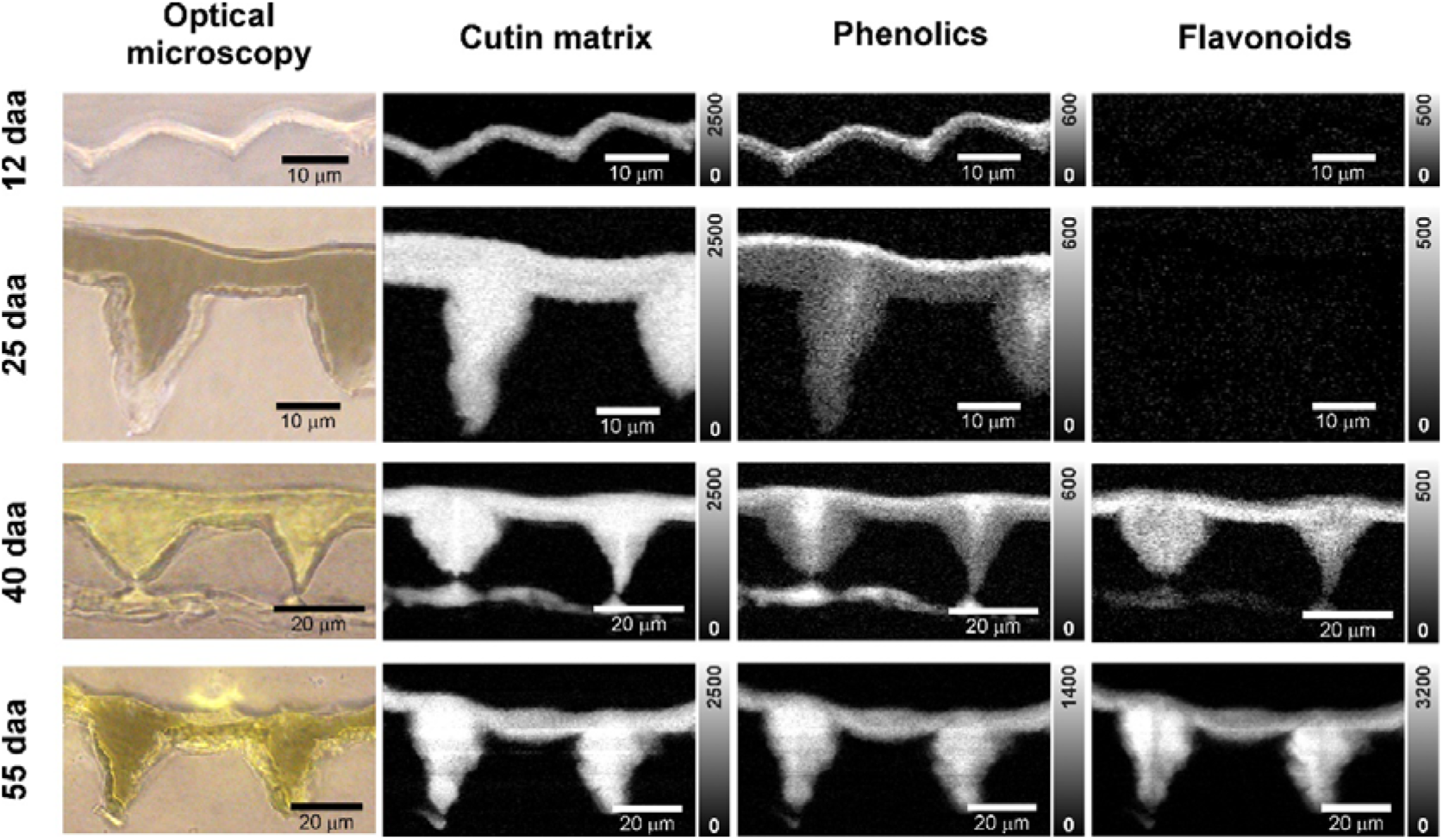
Optical microscopy and Raman images of tomato cuticle cross-sections at different stages of development based on univariate data analysis. ‘Cascada’ cuticles at 12, 25, 40 and 55 daa (days after anthesis) are shown. Raman images are based on the integration of the 1421-1471 cm^−1^ band area, assigned to a cutin enriched matrix, 1622-1642 cm^−1^ band (1618-1638 cm^−1^ in 55 daa due to band shift), mainly attributed to phenolic acids and 1536-1566 cm^−1^ band, corresponding to flavonoids. Colour scale in Raman images is expressed in CCD cts (charge-coupled device counts); note different scaling of phenolic acids and flavonoids in the ripe stage (55 daa).

Phenolic acids impregnate the cuticle already at 12 and 25 daa and accumulate in the outermost layer; most pronounced at 25 daa. Phenolic compounds have been reported in tomato fruit epicuticular waxes (Hunt and Baker, 1980). Thus, to determine the contribution of epicuticular waxes to the phenolic layer identified in this outermost region of the cuticle, the phenolic imaging was repeated after epicuticular wax removal. As the phenolic band neither disappeared nor was reduced after epicuticular wax extraction (Fig. S2), it is evident that phenolics are embedded within the cutin matrix. In later developmental stages, phenolic acids were also observed in the pegs, especially in their centers, which correspond to the outermost part of anticlinal walls and/or middle lamella. At red ripe stage, a higher band area indicates a higher accumulation in the pegs than on the outer periclinal section of the cuticle. Flavonoid accumulation showed a different picture than phenolic acid impregnation. It started later during ripening and was more pronounced at the innermost section of the cuticle at breaker stage (40 daa). At red ripe (55 daa) a dramatic increase in flavonoids was detected in the pegs. However, flavonoids showed higher intensities in the areas where, prior to ripening, phenolic acids showed low intensity, suggesting that mainly flavonoids impregnated these regions towards the epidermal layers.

Integration of three Raman bands characteristics of waxes (1060, 1132 and 1292 cm^−1^, Table 1) allowed to identify a very thin superficial wax layer in several stages of development (Fig. S3). Considering its outermost location and spectral wax characteristics, it corresponds to the epicuticular wax layer. The layer seemed to colocalize with the cutin enriched matrix, but its thickness was close or even smaller than the resolution limit and hence this colocalization should be taken with caution.

### Detailed insights into microchemistry by multivariate approaches

To overcome the problem of Raman band overlaps in the band integration approach and to verify spectra and image interpretation, multivariate data analysis approaches were applied. First, the purest component spectra were identified at different stages of development by Non-negative Matrix Factorization (NMF, Fig.4A-F) and the interpretation of these basis spectra verified by a mixture analysis approach (Fig. 4G-I). These purest basis spectra were used as representative component spectra in a least square fit at every pixel (basis analysis) throughout all developmental stages for comparative imaging (Fig.S4). Finally in-depth mixture image analysis revealed specific components, including different aromatic components and carbohydrates by an orthogonal matching pursuit algorithm (Fig. 5).

**Figure 4.**
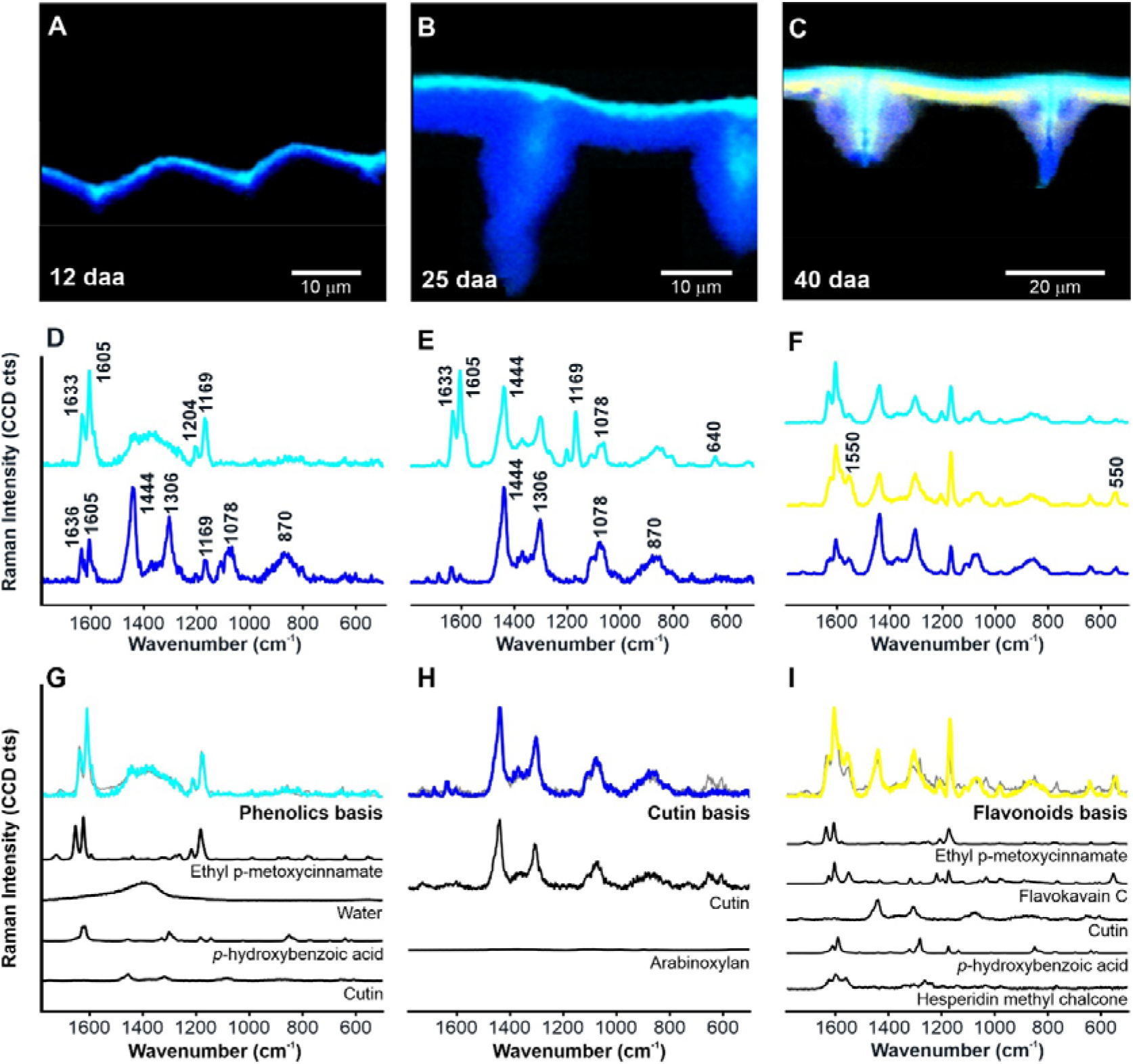
Component distribution maps and the corresponding basis spectra from unmixing analysis (non-negative matrix factorization, NMF) of cuticles from selected stages of development. Basis spectra were assigned using a mixture analysis (orthogonal matching pursuit, OMP) based on a reference library. Component distribution images of ‘Cascada’ fruit cuticle cross-sections at **(A)** 12, **(B)** 25 and **(C)** 40 daa (days after anthesis) based on selected basis spectra **(D-F)** from NMF analysis. Turquoise represents spectra rich in phenolic acids, whereby the purest phenolic acid spectrum was found at 12 daa (**D**). Blue represents cuticle matrix spectra, with a contribution of phenolic acids at 12 daa (**D**), but an almost pure cuticle matrix spectrum at 25 daa (**E**). The yellow spectrum is enriched in flavonoids but with a contribution of phenolic acids and cuticle matrix (**F**). **(G-I)** Mixture analysis of the three selected basis spectra to verify the purity and the composition based on reference spectra. Model fit spectra based on OMP are shown in grey. The reference spectra chosen to fit the basis spectra are shown below. A black canvas was added to the cross-sections for aesthetic reasons.

**Figure 5.**
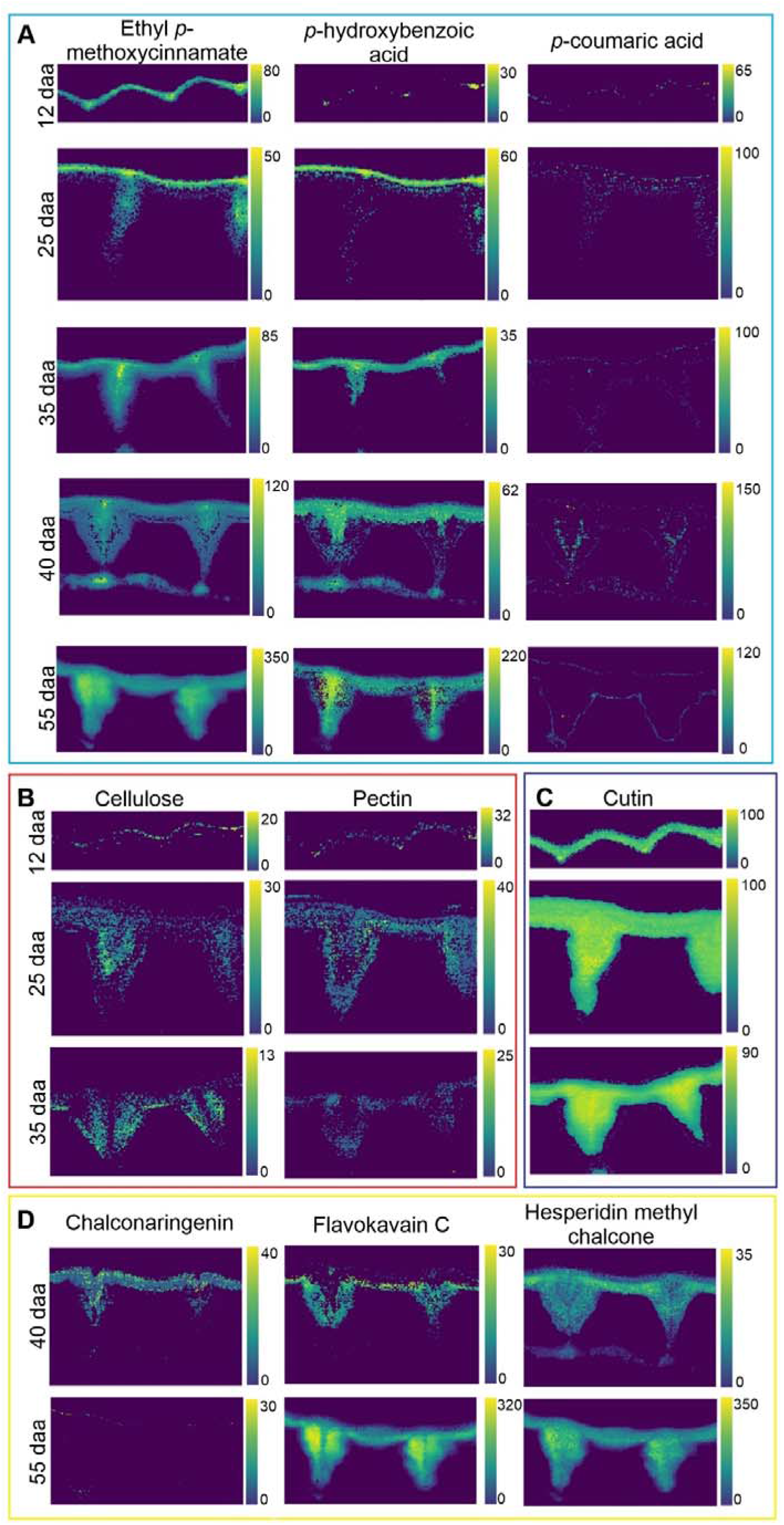
Mixture analysis unravels details on aromatics, carbohydrates and cutin in the tomato fruit cuticle at different stages of development. **(A)** Among phenolic acids, *p*-hydroxybenzoic acid and an ester of *p*-coumaric acid are co-located in a top layer and in the middle of the pegs at unripe stages, while at ripe stages the ester of *p*-coumaric acid becomes more homogenously distributed. **(B)** Cellulose and pectin are found in the unripe state towards the epidermis, while **(C)** cutin displays a homogenous pattern through the cuticle. **(D)** In the ripe stages three different flavonoids are suggested by the algorithm at the inner side of the cuticle. Chemical structures of the components are shown in Fig.S7.

#### Finding the purest cuticle component spectra by NMF

Due to the cuticlés multicomponent nature, the average spectra of the cuticle include bands from all the components (Fig.1). Imaging with a high spatial resolution and at different developmental stages opens the chance to have at least at some pixels at certain stages the one or the other component solely present. Among the different stages, the purest components have been revealed in 12 daa (phenolic acids), 25daa (cutin) and 40 daa (flavonoids) (Fig. 4A-F), whereas others mainly resulted in mixed basis components (Fig. S5). In the very early stage at 12 daa, Raman spectra decomposition revealed two main basis component spectra (Fig. 4D). The first one corresponded to almost pure phenolic acids (turquoise in Fig 4D) with the characteristic bands at 1633, 1605 and 1169 cm^−1^. This “pure” phenolic acid spectrum was only present in the outer part of the cuticle. The second calculated component (blue in Fig. 4D) mainly corresponded to cutin, but also showed a band around 1078 cm^−1^ that could be assigned to either polysaccharides or waxes. Hence, this basis spectrum represented the cuticle matrix (cutin + polysaccharides and/or waxes), but still included a small contribution of phenolic acids. At 25 daa also two main components were distinguished (Fig. 4E): one corresponding to a rich phenolic acids fraction (turquoise in Fig. 4E) but with bands associated to the cuticle matrix, whereas the second one (blue in Fig. 4E) corresponded to an almost pure cuticle matrix spectrum with little to none contribution of phenolic acids. NMF results at 40 daa (Fig. 4F) revealed no “pure” component spectra, but comprised bands assigned to different components and thus the spectra represent mixtures of cuticle components. One is located on top and is enriched in phenolic acids with a small contribution of cuticle matrix (turquoise in Fig. 4F). The layer underneath (yellow in Fig. 4F) corresponds to flavonoids with contributions of phenolic acids and cuticle matrix. The third component (blue in Fig. 4F) was located to the innermost side of the cuticle and showed bands of the cuticle matrix, with little contribution of phenolic acids and flavonoids. None of them corresponded to a pure flavonoid spectrum, although this was the first stage the flavonoids were detected. To derive a purer flavonoid spectrum for basis analysis, we therefore subtracted the cuticle matrix signals from the yellow spectrum (Fig. 4F) (see Materials and Methods for details).

#### Verifying the purity and spectral signature of the basis spectra by Orthogonal Matching Pursuit (OMP) and reverse linear combination of spectra for component imaging

The purity and composition of the basis spectra for phenolic acids and cuticle matrix from 12 and 25 daa, respectively, and the flavonoid spectrum from 40 daa were assessed by modelling them as linear combination of spectra acquired from reference compounds (Fig. 4G-I) by orthogonal matching pursuit (OMP) (Bock et al., 2021). This analysis confirmed the three selected basis spectra as representatives of the three components: cutin, phenolic acids and flavonoids (Fig. 4G-I). The representative cutin spectrum (blue in Fig. 4E) was confirmed as cutin with small contribution of polysaccharides (Fig.4H), while the other two (Fig. 4 G,I) were elucidated as mixtures of different aromatics and still including small contributions of cutin and water. Yet, the basis, selected from 12 daa, (turquoise spectrum in Fig. 4D), was dominated by phenolic acids and esters (Ethyl *p*-methoxycinnamate, *p*-hydroxybenzoic acid, see Figure 4G) and the one selected from 40 daa (yellow spectrum in Fig. 4F) detected in a similar extent phenolics, cutin and flavonoids (hesperidin methyl chalcone and flavokavain C, see Figure 4I). The spectrum related to the presence of flavonoids (yellow in Fig. 4F) was also assessed after subtracting the cutin matrix (Fig.S6), confirming the dominance of flavonoids in this basis spectrum.

In the next step the verified purest component spectra were used to analyse the hyperspectral datasets of all developmental stages during the whole period of fruit growth and ripening. The three selected component spectra and a water spectrum were linearly combined at each pixel using a basis analysis to finally result in distribution maps of all four components (Fig.S4). The cuticle matrix (cutin + polysaccharides and/or waxes) was homogenously distributed throughout the cuticle in all developmental stages (blue in Fig.S4), whereas for phenolic acids (turquoise in Fig.S4) and flavonoids (yellow in Fig.S4) we confirmed the spatial and temporal changes as detected by band integration (Fig.3), but with more details (Figure S4). Additionally water was included in the fit (purple in Fig.S4) and revealed to accumulate nearby the inner side of the cuticle. The flavonoid component spectrum was not present in the datasets of earlier stages, but appeared in traces with the onset of ripening at 35 daa. At 40 daa these compounds were clearly located underneath the phenolic acid layer and across the entire width of the pegs (see Fig. S4). With fruit ripening, flavonoids continued to accumulate across to the entire cuticle. At red ripe, a contrasting pattern of phenolic acids and flavonoids was typical: phenolic acids close to the outer region and in the central region of the pegs, whereas the flavonoid signal more intense underneath this phenolic acid layer and across the entire pegs.

#### In-depth hyperspectral dataset analysis by OMP to reveal specific components

As the multicomponent nature of the cuticle was proven through all developmental stages and even on the microscale of 333 nm (pixel size) (Fig. 4 G-I), we applied the mixture analysis on the hyperspectral datasets. At every pixel, a model fit is searched for by the OMP algorithm based on our reference spectral dataset, including carbohydrates, aromatics and lipids (Fig. 5). This analysis included the novelty of selectively locating the most relevant phenolics along with carbohydrates and cutin.

Among the phenolics detected throughout the development was *p*-coumaric acid mainly in esterified form and *p*-hydroxybenzoic acid (Fig. 5 A). The two were colocated in a superficial layer and in the middle area of the pegs in the early stages of development, but the ester became more homogenous in the ripe stage (Fig. 5A). In the early unripe stages we could moreover locate carbohydrates (pectin and cellulose) (Fig. 5B) as well as cutin (Fig. 5C). While the latter one showed again a very homogenous distribution throughout the whole cuticle (Fig. 5C), the cellulose and pectin accumulated in contrary areas than the phenolic acids (Fig. 5B). However, the carbohydrate rich regions towards the epidermis seem to coincide with flavonoid locations during ripening (Fig. 5D). Among the flavonoids, chalconaringenin was only detected in small with small parameter estimates at 40 daa (Fig.5D), whereas at most pixels the cuticle spectra were better represented by methoxylated chalcones, such as flavokavain C and hesperidin methyl chalcone (see structures in Fig. S7 and their Raman spectra in Fig. 4I). However, at the ripe state the error of the fit increased and several observed band mismatches were observed. Although numerous flavonoids and aromatic were included in the library, it became more and more difficult to fit the pixel spectra by the reference spectra and thus pointed to more interactions in the mature stages.

## Discussion

Using Confocal Raman Microscopy we successfully monitored topochemical changes of lipids, carbohydrates and aromatic components during tomato fruit cuticle development. By combining uni-and multivariate analysis we assigned and verified Raman bands and/or spectra to the main component groups and analyzed their spatial distribution as well as their changes from green to red stage (Fig. 1-4). Especially, the mixture analysis combined with a comprehensive spectral reference library has provided detailed insights into the microdistribution of cutin, cellulose, pectin and the diversity of aromatic molecules (Fig. 5). Finally, all new insights were included in 3 schematic models to summarize our understanding of the tomato fruit cuticle and its development (Fig. 6).

**Figure 6.**
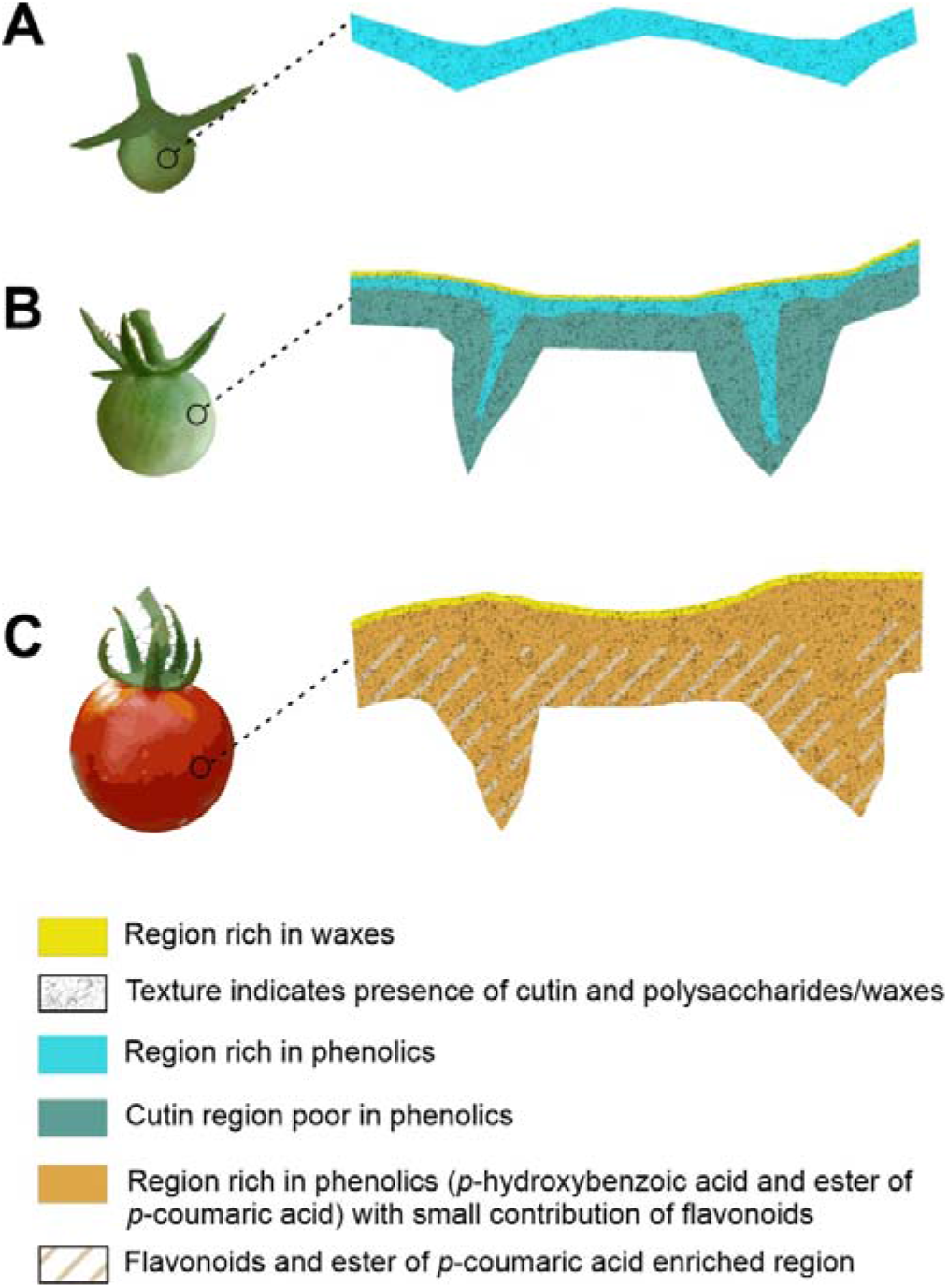
Schematic representation of the spatial location of the main cuticle components throughout tomato fruit development. **(A)** the earliest stages of development (8-12 daa) showed a cuticle matrix enriched in phenolic acids (cyan). **(B)** During most of the growing period (15-35 daa) phenolic acids were only located at the outer region of the cuticle and above a very thin superficial waxy layer was detected **(C)** During ripening, *p*-hydroxybenzoic acid preserves its initial location (cyan) while the ester of *p*-coumaric acid extend throughout the entire cuticle width (orange) and flavonoids showed a gradient with higher deposition towards the inner side of the cuticle (stripes). A cuticle matrix (cutin plus polysaccharides and/or waxes) is present across the entire cuticle width in all the stages of development (texture). daa, days after anthesis.

### Cutin and polysaccharides: gradual and/or uniform foundation of the cuticle?

A heterogeneous distribution of the two matrix polymers has been reported. Cutin is regarded to decrease towards the inner side of the cuticle, while at the same time polysaccharides increase (Reynoud et al., 2021) (Fernández et al., 2016). Yet, Raman bands assigned to the cuticle matrix (Fig.3), and also the model fits with cutin spectra (Fig.5, Fig. S4) revealed no cutin intensity gradient in any of the stages of development. This is an indication that the amount of cutin is quite uniform within the tomato fruit cuticle. Waxes are embedded as intracuticular waxes and were additionally imaged as a very thin superficial layer (Fig. S3). The wax specific bands are assigned to long-chain alkanes or alcohols, compounds that have been reported in the epi-and intracuticular wax fractions of tomato fruit (Prinsloo et al., 2004; Vogg et al., 2004).

No strong Raman marker bands were found for carbohydrates; not for cellulose/hemicellulose and not for pectin and not at any pixel throughout all stages. As the Raman intensity is not only a matter of the amount of the molecules present, but also depends on the polarizability of the molecules, weaker Raman scatterers, like carbohydrates, can get overlaid by lipids and aromatic components. This is the reason why carbohydrates were only detected in the younger developmental stages (Fig. 5B), although they are reported to make up almost one third of the cuticle (25-30%) (Domínguez et al., 2008). Due to the small contribution of carbohydrates to the overall cuticle Raman signal, no strong marker bands were detected and their distribution only elucidated with the help of mixture analysis (Fig. 5B). With this approach we found cellulose and pectin in the younger stages especially in the pegs and towards the epidermal layer, agreeing with their previously reported location along cuticle transversal sections of some tree species (Guzmán et al., 2014a; Guzmán et al., 2014b). In the mature stage, the higher background and signal of flavonoids masks the cutin bands partly and the carbohydrates completely.

### Tomato fruit procuticle is enriched in phenolic acids

Although phenolic components are reported to be around 0.8-7.5% depending on the stage of development in the tomato fruit cuticle (España et al., 2014b), they are easily detected by Raman imaging due to their strong Raman scattering intensity resulting in numerous strong bands.

During the early stages of development, phenolics (mainly esterified *p*-coumaric acid at 12 daa according to Fig.5) can be located throughout the overall thickness of the procuticle. This initial and uniform procuticle is present in tomato fruit during the cell division period (Segado et al., 2016). Indeed, a small decrease in the phenolics/cutin ratio can be observed later on during the beginning of the cell expansion period (Fig. 2B) that agrees with the reduction in percentage of phenolics detected during the 15-30 daa period (~0.8%) compared to the previous stage (2.5% at 10 daa) (data from (España et al., 2014b)). A similar percentage of phenolics was only achieved and exceeded during ripening. The increase in cuticle deposition during the expansion period can then be attributed to cutin with little participation of phenolics, thus decreasing the phenolics/cutin ratio. The notable accumulation of phenolics in the outer cuticle region remained during growth and exceeded also in the outermost layer of the cutinized anticlinal cell walls, that is the middle region of the so-called pegs. These phenolics were identified as a mixture of esterified *p*-coumaric acid and *p*-hydroxybenzoic acid. In several species, cinnamic acid derivatives have been identified esterified to the hydroxyl fatty acids of cutin (Cheng et al., 2013; Renault et al., 2017; Reynoud et al., 2021). *In situ* identification of *p*-coumaric acid as ethyl *p*-methoxycinnamate in the basis spectra and in the mixture analysis of different stages of development of tomato fruit cuticle supports the hypothesis of this molecule acting either as a non-lipid cutin monomer or as a link between the polysaccharide and cutin domains, esterified by both the acid and the hydroxyl groups.

We detected phenolic acids already in the earliest stages of development in the procuticle in high amounts. A fact, also interesting under the aspect that early land plants like different bryophytes have a high percentage of phenolics (35%) and thus a phenolic enriched cutin (Renault et al., 2017; Kong et al., 2020). This posits the idea that a high percentage of cuticle phenolics could not necessarily be associated with a specific plant group but to the initial procuticle stage and to the degree of cuticle deposition in later stages of development. Thus, in species where the cuticle would not further increase its deposition past the initial procuticle stage, or have a limited increase, a phenolic enriched cuticle could be detected, whereas in those species with notable cuticle increase after the procuticle deposition, the final result would be a cuticle with a lower percentage of phenolics. This hypothesis deserves further study as well as the link observed in tomato fruit between the procuticle and the cell division stage of growth and the cell expansion period with a massive cuticle deposition (Segado et al., 2020).

A remarkable degree of UV protection achieved by the cuticle has been reported and discussed in context with phenolic acids (Krauss et al., 1997; Benítez et al., 2022; González Moreno et al., 2022). The tomato fruit cuticle was shown to block 95% of the incident UV-B light at the early stages and reaching 99% later in development, whereas in other species and organs, these results varied from an almost complete blockage to over 50%. Cinnamic acid derivatives, such as *p*-coumaric acid present in tomato fruit cuticle and in all the studied cuticles, considerably absorb UV-B light. Hence, these molecules give the cuticle special optical properties to protect plant tissues from harmful UV light. Thus, our identified outer layer of phenolic acids in tomato fruit cuticle throughout the development represents an optimal location to block the UV-B already at entry.

### Phenolic acids and flavonoids accumulate during tomato fruit ripening

Also during ripening *p*-coumaric and *p*-hydroxybenzoic acids increase in the tomato fruit cuticle (España et al., 2014b). This is in agreement with the CRM results of the semi-quantitative mixture analysis, which showed about four times higher contributions of phenolic acids in the mature state than in the green state (Fig.5A). Also in the mature stage the OMP suggested ethyl trans-4-methoxycinnamate in esterified form and the free *p*-coumaric acid only at a few positions and to less extent (Fig. 5A, Fig. 4G, I). These results suggest that the huge majority of *p*-coumaric acid is present within the cuticle as an ester. It coexists during the whole development with the other main phenolic acid reported in tomato fruit cuticle, the *p*-hydroxybenzoic acid.

However, the dramatic increase in cuticle phenolics during tomato ripening has been mainly ascribed to *de novo* synthesis of the flavonoid chalconaringenin (España et al., 2014a). This was proven by a 60-70% reduction of the cuticle phenolics through transient silencing of *CHALCONE SYNTHASE*, which is responsible for chalconaringenin synthesis (España et al., 2014a). The expression of the next gene of the flavonoid pathway, *CHALCONE ISOMERASE* (*CHI*), is hardly detected during tomato fruit ripening (Bovy et al., 2002), hence mainly chalconaringenin is accumulated in the cuticle of ripe fruits (Hunt and Baker, 1980). Chemical isomerization between chalconaringenin and naringenin flavone (the product of CHI) has been reported (Kaltenbach et al., 2018), but recent quantum chemical analysis has shown that this chemical isomerization is very unlikely within the cuticle matrix (González Moreno et al., 2021). Raman imaging together with univariate and multivariate analysis confirmed a higher amount of flavonoids in the cuticle during ripening (Figure 1-6). However, chalconaringenin was only selected by the OMP algorithm at 40 daa (Fig. 5), but not at ripe stages. The mismatch of the cuticle spectra and the chalconaringenin spectrum was also evident in direct comparison of the spectra. When we added also a methoxylated chalconaringenin and a methoxylated and glycosylated chalcone to the library, the OMP algorithm chose the two and the model fit explained the cuticle spectra better (see structures in Fig. S7 and their Raman spectra in Fig. 4G-I). Flavonoids located on the surface of plants have been mostly identified as aglycones, often with substituted methoxy groups (Onyilagha and Grotewold, 2004), a modification that increases molecule hydrophobicity. Hence, it is possible that the tomato fruit cuticle accumulates some methoxylated modification of chalconaringenin. But as the spectra of the ripe tomato did not fit the references perfectly, changes in the environment and even interaction with other cuticle components could be the reason behind the band differences in Raman spectra observed of pure chalconaringenin and that present in the cuticle of red ripe fruits. This means that chalconaringenin is either methoxylated, glycosilated or covalently bond to other cuticle components over its A-ring OH-groups (at C4 and C6; the OH on C2 hydrogen-bonds to the carbonyl oxygen).

Impregnation with cinnamic acid derivatives and flavonoids confers mechanical resistance and structural rigidity to the cutin matrix during ripening (Benítez et al., 2021; Benítez et al., 2022). Thus, the reported gradual changes in thermal and mechanical properties of the tomato fruit cuticle during ripening are mirrored in our results by the increasing deposition of these phenolics. At the ripe stage the ester of *p*-coumaric acid was present throughout the entire width of the cuticle and pegs (Fig. 5A). Flavonoids were also distributed throughout the whole cuticle with highest amounts towards the inner side of the cuticle (Fig. 5D). The distribution pattern across the entire cuticle supports their above mentioned structural role as fillers that restrict the molecular motion of cutin chains. Our results revealed the carbohydrates in the earlier stages at locations (Fig. 5B) where the flavonoids start to impregnate the cuticle during ripening (Fig. 5C). Thus, flavonoids may finally fill the spaces within the more hydrophobic carbohydrate network and link them to the hydrophilic cutin polymer.

### Three schematic models depict microchemistry of tomato fruit cuticles during development

Based on Raman imaging of all cuticle components, we distinguish three different models during tomato cuticle development (Fig 6). The first one corresponds to the procuticle stage in the very beginning, where phenolic acids (mainly as esterified *p*-coumaric acid) are present throughout the entire cuticle matrix. The second distribution model was observed from 15 (beginning of the cell expansion period) to 35 daa (mature green stage), in the period with most fruit growth. In this stage, the cuticle showed a thin superficial waxy layer. This layer colocalized with the cuticle matrix (cutin with polysaccharides and waxes), an indication that it could be embedded within the cuticle. It was not readily identified in all the stages of development, most probably due to its thickness being close or lower to the resolution limit of CRM. Underneath the wax layer, a phenolic acid enriched cuticle layer was observed. This phenolic layer was not affected by epicuticular wax removal indicating that, at least, most of these phenolic acids are integral part of the cuticle matrix. This phenolic acid layer was also located to the outermost region of the cutinized anticlinal cell walls. Beneath the phenolic layer, a much thicker layer corresponding to a cuticle matrix without phenolics extended to the inner side of the cuticle. Finally, during ripening, additional incorporation of phenolic acids and the deposition of flavonoids takes place. *p-*Hydroxybenzoic acid preserves the initial location of phenolics within the cuticle; but, *p*-coumaric acid (mainly as an ester) is incorporated to the inner cuticle matrix layer, which was devoid of phenolics, thus covering the entire cuticle thickness and peg width. On the other hand, flavonoids show a gradient across the cuticle, with higher concentration towards the inner side of the cuticle, in closer contact with the epidermal cell wall, and within pegs outside the central region.

To summarize, the high Raman scattering intensity of aromatic components gives new insights into the incorporation of phenolic acids and flavonoids during development. The dual location of phenolic acids identified during growth and later during ripening could be associated to different biological functions. Phenolic acids might act primarily as an outer photoprotective layer during growth and contribute to mechanical stiffening and strengthening of the cuticle during ripening, together with flavonoids. The cutin matrix showed a similar intensity across the cuticle and development, while carbohydrates were due to their low Raman scattering only detected in early stages. Accumulation at regions, where later flavonoids start to impregnate the cuticle, suggest filling of hydrophilic regions with aromatics. Spatial co-locations as well as spectral misfits with chalconaringenin point to interconnections of the flavonoids with other cuticle components.

## Materials and Methods

### Sample preparation and micro-sectioning

Tomato fruit cuticles of different stages of development, from 8 days after anthesis (daa) to 55 daa (ripe red), of *Solanum lycopersicum* L. ‘Cascada’ were isolated using an enzymatic protocol (Petracek and Bukovac, 1995; España et al., 2014b). Samples were cross-sectioned into slices of 10 μm thickness by cryo-microtomy at −15°C (CM3050 S, Leica, Austria). The transversal sections were put onto microscope slides with a drop of D_2_O. They were covered with a cover slip of 0.17 mm thickness and sealed with nail polish to prevent water evaporation during measurement.

### Confocal Raman Microscopy

Raman spectra from isolated tomato cuticles were acquired by using confocal Raman microscopy (alpha300RA, WITec GmbH, Germany). Samples were focused with a 100x oil immersion objective (numerical aperture (NA)=1.4, correction of the coverslip of 0.17 mm). A linear polarized coherent diode laser λ_ex_ = 785 nm (XTRAII, Toptica Photonics, Germany) was used as excitation line. The Raman signal was received by an optic multifiber (diameter of 100 μm), transported into the spectrometer UHTS 300 (WITec, Germany; 600g.mm^−1^ grating, spectral resolution around 3.8 cm^−1^) and intensities measured using a CCD camera DU401A BR DD (Andor, Belfast, North Ireland). The laser power was set at 150 mW and the integration time to 0.1 s. Spatial resolution of the measurement was calculated by r= 0.61λ/NA, being r, λ and NA the lateral resolution, excitation laser wavelength and numerical aperture of the objective, respectively. Maximum spatial resolution achieved was around 342 nm. The size of evaluated scan areas was from 20 μm x 10 μm (200 μm^2^) to 85 μm x 60 μm (5100 μm^2^). A minimum of three cross-sections from three different fruits of each stage of development have been measured to obtain robust results.

### Raman band assignments

The assignment of some characteristic vibrational bands of main compounds in tomato fruit cuticle was done according to literature (Table 1) and *DFT (Density-Functional Theory)* calculations in case of chalconaringenin (predominant flavonoid extracted and characterized in tomato fruit cuticle) due to some bands are not described in literature (see Table S1 for more details). It is worth clarifying that some wavenumbers from the literature are from infrared rather than Raman spectroscopy. Although both techniques differ in the way they excite molecular vibrations, their respective band positions are equivalent.

*DFT* calculations were performed using Gaussian 16 (Frisch et al., 2016) software. Raman spectra were calculated at B3LYP with 6-31+G(d,p) basis set, applying a scale factor of 0.9613 to overcome experimental–theoretical deviations. Band assignments were carried out by *GaussView* software.

### Spectral pre-processing and univariate approaches

Cosmic rays, which are very intense signals generated by the interaction of energetic particles with the CCD camera, were removed before processing acquired data using the software *Project Four 4*.*1* (WITec, Germany). Pixels which show characteristic Raman bands of cuticle were masked in each replicate and the average spectrum of each stage of development was calculated (throughout their three replicates) using *Project Four 4*.*1*. No normalization process was applied to these data in order to show the most realistic comparative overview. Rayleigh lines were removed, and the spectral range limited to 300-1700 cm^−1^ in OPUS (Bruker, Germany). Spectra were integrated at characteristic bands using the same software. At 8daa, chemical imaging of the sample was not possible due to the extreme thinness of the cuticle; at 10 daa an improvement at signal/noise ratio was required to obtain a well-defined spectrum. For this reason, the ratio of pixels recovered per μm of sample was changed from 60 px/20 μm used for all the rest of the samples to 100 px/20 μm.

Further spectral pre-processing actions before band integration, NMF and mixture analysis were smoothing and background subtraction using *Project Four 4*.*1*. The integration of specific bands (see Table 1) of each cuticular component (cutin, polysaccharides, waxes, phenolic compounds and flavonoids), allowed to locate them along the depth of the cuticle.

### Unmixing algorithm: Nonnegative Matrix Factorization (NMF)

As band integration is focusing to individual characteristic peaks only, it shows limitations due to overlapping bands in the spectrum. Spectral unmixing aims to separate measured datasets (D), such as Raman maps, into pure components (S^T^, also termed endmembers or basis) and their respective coefficients (C). Using this method in chemometrics, the pure components often refer to different chemical compounds while the coefficients represent concentration profiles or abundance maps of the respective compounds (van Stokkum et al., 2009). Using mathematical notation, this basic problem can be stated as follows:

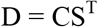

There are several spectral unmixing methods available, such as nonnegative matrix factorization (NMF), vertex component analysis (VCA) or multivariate curve resolution alternating least squares (MCR-ALS), which differ in their optimization approach and constraints (Prats-Mateu et al., 2018b).

Nonnegative Matrix Factorization (Lee and Seung, 1999) might result in a more realistic representation of the spatial abundance of different compounds than univariate approaches. Runs with different numbers of basis from 2 to 5 were done and, after evaluating the pure basis spectra, the most meaningful result kept. The maximum number of iterations was set to 10.000. From the basis spectra of all NMF runs among the different images, a reference library representing cutin, phenolics, flavonoids and water to be used in basis analysis was compiled. To achieve a more selective flavonoid location, consecutive subtractions of cutin-polysaccharides signature were applied to flavonoid spectrum until reach the optimal coefficient (0.35) which minimizes the presence of the cuticle matrix but without observing any negative band along the spectra (pink spectrum in Figure S6).

The interpretation of the obtained basis was confirmed using measured spectra from reference compounds (Fig.4G-I). To do so, an orthogonal matching pursuit (Pati et al., 1993) was used to select predictors to be used in a linear model to approximate the respective basis spectrum.

### Basis analysis

Basis analysis was used to acquire the linear combination of the Raman spectra of the basis (obtained by NMF) which best represents the bidimensional spectrum of each pixel of the sample. These estimated coefficients are calculated by a least square fit, and are proportional to the abundance of each compound in the mixture (Schmidt et al., 2005). As each pixel can be separately analysed, the application of basis analysis in confocal Raman measurements allows to assess the distribution of compounds in space getting colour maps for each component.

### Spectral reference library and mixture analysis

Isolated tomato fruit cutin, regardless the stage of development, accumulates phenolic compounds (España et al., 2014b). Thus, to obtain a reference spectrum of cutin, a recombinant inbred line (RIL115) derived from a cross between the domesticated tomato and a wild species (Alba et al., 2009) was employed. This line has the characteristic of having virtually zero phenolic compounds in its cuticle (Domínguez et al., 2009a) thanks to a specific combination of a number of genomic regions related to the accumulation of phenolic compounds in the cuticle (Barraj Barraj et al., 2021). Beside the cutin reference the library included 148 entities, including water, aromatics, lipids, minerals and carbohydrates.

To analyze the multicomponent nature of the cuticle spectra, we modelled them as a linear combination of measured reference spectra using the Orthogonal Matching Pursuit (OMP) (Pati et al., 1993) The OMP is an iterative and fast approach that can be applied to spectra and spectral images. At each iteration it seeks for a component of the spectral reference library, which is best correlated with the residual of the linear combination of references selected in the preceding iterations (the experimental spectrum, being the residual from an empty model, used in the first iteration) and adds it to the predictor set. A stopping criterion (e.g., a threshold for residual error or a maximal number of iterations) stops the search for additional members. The predictor set of the final model holds the selected constituents of the mixture, and the coefficients indicate the abundance of the respective compounds.

## Supplemental Data

**Table S1**. Raman band assignment of chalconaringenin in gas phase according to DFT calculations.

**Figure S1**. Raman spectra of the free conformations of *p*-coumaric acid and chalconaringenin.

**Figure S2. Raman images corresponding to a univariate data analysis of phenolic acids in tomato cuticle cross-sections before (left) and after (right) epicuticular wax removal**. Raman images of ‘Cascada’ fruit cuticles at 15 daa (days after anthesis) before (left) and after (right) extraction of epicuticular waxes. These images have been generated for the integration of the 1622-1642 cm-1 band mainly attributed to phenolic acids. Images were scaled from minimum (black) to maximum (bright colours).

**Figure S3. Raman images based on integration of a band related to the presence of a superficial waxy layer**. Raman images were generated by integrating over 1127-1137 cm^−1^ (waxes) of ‘Cascada’ fruit cuticles from different stages of fruit development. The grey scale of the Raman images a represents CCDcts (Charge Coupled Device counts).

**Figure S4. Basis analysis using the purest spectra obtained from the NMF algorithm**. Distribution maps after a basis analysis of different cuticle components in cross-sections of ‘Cascada’ fruit cuticles throughout development, from 10 to 55 daa (days after anthesis). Location of phenolics acids, flavonoids with a contribution of phenolic acids, the cuticle matrix (cutin + polysaccharides + waxes) and water are shown in turquoise, yellow, blue and purple, respectively. False colours within each image were scaled to minimum (black) to maximum (bright colours).

**Figure S5. Distribution maps and Raman spectra obtained by NMF at 10 daa and 55 daa**. Abundance maps of ‘Cascada’ fruit cuticles at 10 and 55 daa. They are accompanied by extracted spectra of each basis as obtained by NMF. Turquoise, blue and yellow colours represent the basis with dominant contribution of phenolics, cuticle matrix and flavonoids, respectively. A black canvas was added to the cross-sections for aesthetic reasons.

**Figure S6. Mixture analysis of the basis spectra of the flavonoid spectra after subtracting cuticle matrix contribution (pink spectrum)**. The basis spectrum fit solely with phenolics and flavonoids from the library. Model fit spectra based on OMP are shown in grey.

**Figure S7**. Chemical structure of **(A)** *p*-hydroxybenzoic acid **(B)** *p*-coumaric acid **(C)** ethyl *p*-methoxycinnamate **(D)** chalconaringenin **(E)** flavokavain C **(F)** hesperidin methyl chalcone.

## Acknowledgments

The authors thank the Raman measurements technical advice and support from Nadia Sasani and Martin Felhofer (Institute of Biophysics, University of Natural Resources and Life Sciences, Vienna, Austria).

## References

Alba JM, Montserrat M, Fernández-Muñoz R (2009) Resistance to the two-spotted spider mite (Tetranychus urticae) by acylsucroses of wild tomato (Solanum pimpinellifolium) trichomes studied in a recombinant inbred line population. Exp Appl Acarol 47: 35–47

Barraj Barraj R, Segado P, Moreno-González R, Heredia A, Fernández-Muñoz R, Domínguez E (2021) Genome-wide QTL analysis of tomato fruit cuticle deposition and composition. Hortic Res 8: 113

Benítez JJ, González Moreno A, Guzmán-Puyol S, Heredia-Guerrero JA, Heredia A, Domínguez E (2022) The response of tomato fruit cuticles against heat and light. Front Plant Sci 12: 807723

Benítez JJ, Guzmán-Puyol S, Vilaplana F, Heredia-Guerrero JA, Domínguez E, Heredia A (2021) Mechanical performances of isolated cuticles along tomato fruit growth and ripening. Front Plant Sci 12: 787839

Bock P, Felhofer M, Mayer K, Gierlinger N (2021) A Guide to Elucidate the Hidden Multicomponent Layered Structure of Plant Cuticles by Raman Imaging. Front Plant Sci 12: 793330

Bovy A, De Vos R, Kemper M, Schijlen E, Almenar Pertejo M, Muir S, Collins G, Robinson S, Verhoeyen M, Hughes S, et al (2002) High-flavonol tomatoes resulting from the heterologous expression of the maize transcription factor genes LC and C1. Plant Cell 14: 2509–2526

Casado CG, Heredia A (2001) Specific heat determination of plant barrier lipophilic components: Biological implications. Biochim Biophys Acta - Biomembr 1511: 291–296

Cheng AX, Gou JY, Yu XH, Yang H, Fang X, Chen XY, Liu CJ (2013) Characterization and ectopic expression of a populus hydroxyacid hydroxycinnamoyltransferase. Mol Plant 6: 1889–1903

Domínguez E, España L, López-Casado G, Cuartero J, Heredia A (2009a) Biomechanics of isolated tomato (Solanum lycopersicum) fruit cuticles during ripening: The role of flavonoids. Funct Plant Biol 36: 613–620

Domínguez E, Heredia-Guerrero JA, Heredia A (2011) The biophysical design of plant cuticles: An overview. New Phytol 189: 938–949

Domínguez E, López-Casado G, Cuartero J, Heredia A (2008) Development of fruit cuticle in cherry tomato (Solanum lycopersicum). Funct Plant Biol 35: 403–411

Domínguez E, Luque P, Heredia A (2009b) Sorption and interaction of the flavonoid naringenin on tomato fruit cuticles. J Agric Food Chem 57: 7560–7564

España L, Heredia-Guerrero JA, Reina-Pinto JJ, Fernández-Muñoz R, Heredia A, Domínguez E (2014a) Transient silencing of CHALCONE SYNTHASE during fruit ripening modifies tomato epidermal cells and cuticle properties. Plant Physiol 166: 1371–1386

España L, Heredia-Guerrero JA, Segado P, Benítez JJ, Heredia A, Domínguez E (2014b) Biomechanical properties of the tomato (Solanum lycopersicum) fruit cuticle during development are modulated by changes in the relative amounts of its components. New Phytol 202: 790–802

Farber C, Li J, Hager E, Chemelewski R, Mullet J, Rogachev AY, Kurouski D (2019a) Complementarity of Raman and infrared spectroscopy for structural characterization of plant epicuticular waxes. ACS Omega 4: 3700–3707

Farber C, Wang R, Chemelewski R, Mullet J, Kurouski D (2019b) Nanoscale structural organization of plant epicuticular wax probed by atomic force microscope infrared spectroscopy. Anal Chem 91: 2472–2479

Fernández V, Guzmán-Delgado P, Graça J, Santos S, Gil L (2016) Cuticle structure in relation to chemical composition: Re-assessing the prevailing model. Front Plant Sci 7: 427

Frisch MJ., Trucks GW., Schlegel HB., Scuseria GE., Robb MA., Cheeseman, J. R.; Scalmani G., Barone V., Petersson GA., Nakatsuji, H.; Li X., et al (2016) Gaussian 16, Revision A.03.

Gautam R, Vanga S, Ariese F, Umapathy S (2015) Review of multidimensional data processing approaches for Raman and infrared spectroscopy. EPJ Tech Instrum 2: 1–38

Gierlinger N (2018) Raman imaging of plant cell walls. In O Toporski, J, Dieing, T, Hollricher, ed, Confocal Raman Microsc. Springer Series in Surface Sciences, pp 471–481

Gierlinger N, Schwanninger M (2006) Chemical imaging of poplar wood cell walls by confocal Raman microscopy. Plant Physiol. doi: 10.1104/pp.105.066993

González Moreno A, de Cózar A, Prieto P, Domínguez E, Heredia A (2022) Radiationless mechanism of UV deactivation by cuticle phenolics in plants. Nat Commun 13: 1786

González Moreno A, Prieto P, Ruiz Delgado MC, Domínguez E, Heredia A, De Cózar A (2021) Structure, isomerization and dimerization processes of naringenin flavonoids. Phys Chem Chem Phys 23: 18068–18077

Guzmán P, Fernández V, García ML, Khayet M, Fernández A, Gil L (2014a) Localization of polysaccharides in isolated and intact cuticles of eucalypt, poplar and pear leaves by enzyme-gold labelling. Plant Physiol Biochem 76: 1–6

Guzmán P, Fernández V, Graça J, Cabral V, Kayali N, Khayet M, Gil L (2014b) Chemical and structural analysis of Eucalyptus globulus and E. camaldulensis leaf cuticles: A lipidized cell wall region. Front Plant Sci 5: 481

Hama T, Seki K, Ishibashi A, Miyazaki A, Kouchi A, Watanabe N, Shimoaka T, Hasegawa T (2019) Probing the molecular structure and orientation of the leaf surface of Brassica oleracea L. by polarization modulation-infrared reflection-absorption spectroscopy. Plant Cell Physiol 60: 1567–1580

Heredia-Guerrero JA, Benítez JJ, Domínguez E, Bayer IS, Cingolani R, Athanassiou A, Heredia A (2014) Infrared and Raman spectroscopic features of plant cuticles: A review. Front Plant Sci 5: 305

Heredia A (2003) Biophysical and biochemical characteristics of cutin, a plant barrier biopolymer. Biochim Biophys Acta 1620: 1–7

Hunt GM, Baker EA (1980) Phenolic constituents of tomato fruit cuticles. Phytochemistry 19: 1415–1419

Jetter R, Riederer M (2016) Localization of the transpiration barrier in the epi-and intracuticular waxes of eight plant species: Water transport resistances are associated with fatty acyl rather than alicyclic components. Plant Physiol 170: 921–934

Kaltenbach M, Burke JR, Dindo M, Pabis A, Munsberg FS, Rabin A, Kamerlin SCL, Noel JP, Tawfik DS (2018) Evolution of chalcone isomerase from a noncatalytic ancestor. Nat Chem Biol 2018 146 14: 548–555

Khanal BP, Grimm E, Finger S, Blume A, Knoche M (2013) Intracuticular wax fixes and restricts strain in leaf and fruit cuticles. New Phytol 200: 134–143

Kong L, Liu Y, Zhi P, Wang X, Xu B, Gong Z, Chang C (2020) Origins and evolution of cuticle biosynthetic machinery in land plants. Plant Physiol 184: 1998–2010

Krauss P, Markstädter C, Riederer M (1997) Attenuation of UV radiation by plant cuticles from woody species. Plant, Cell Environ 20: 1079–1085

Lara I, Heredia A, Domínguez E (2019) Shelf life potential and the fruit cuticle: The unexpected player. Front Plant Sci 10: 770

Lee DD, Seung HS (1999) Learning the parts of objects by non-negative matrix factorization. Nature 401: 788–791

López-Casado G, Matas AJ, Domínguez E, Cuartero J, Heredia A (2007) Biomechanics of isolated tomato (Solanum lycopersicum L.) fruit cuticles: The role of the cutin matrix and polysaccharides. J Exp Bot 58: 3875–3883

Mansfield JC, Littlejohn GR, Seymour MP, Lind RJ, Perfect S, Moger J (2013) Label-free chemically specific imaging in planta with stimulated Raman scattering microscopy. Anal Chem 85: 5055–5063

Martin LBB, Rose JKC (2014) There’s more than one way to skin a fruit: Formation and functions of fruit cuticles. J Exp Bot 65: 4639–4651

Mateu BP, Hauser MT, Heredia A, Gierlinger N (2016) Waterproofing in Arabidopsis: Following phenolics and lipids in situ by Confocal Raman Microscopy. Front Chem 4: 10

Mazurek S, Mucciolo A, Humbel BM, Nawrath C (2013) Transmission Fourier transform infrared microspectroscopy allows simultaneous assessment of cutin and cell-wall polysaccharides of Arabidopsis petals. Plant J 74: 880–891

Onyilagha, JC, Grotewold E (2004) The biology and structural distribution of surface flavonoids. Recent Res Dev Plant Sci 2: 53–71

Pati YC, Rezaiifar R, Krishnaprasad PS (1993) Orthogonal matching pursuit: recursive function approximation with applications to wavelet decomposition. Conf. Rec. Asilomar Conf. Signals, Syst. Comput. pp 40–44

Petracek PD, Bukovac MJ (1995) Rheological properties of enzymatically isolated tomato fruit cuticle. Plant Physiol 109: 675–679

Philippe G, Geneix N, Petit J, Guillon F, Sandt C, Rothan C, Lahaye M, Marion D, Bakan B (2020) Assembly of tomato fruit cuticles: a cross-talk between the cutin polyester and cell wall polysaccharides. New Phytol 226: 809–822

Prats-Mateu B, Bock P, Schroffenegger M, Toca-Herrera JL, Gierlinger N (2018a) Following laser induced changes of plant phenylpropanoids by Raman microscopy. Sci Rep 8: 11804

Prats-Mateu B, Felhofer M, de Juan A, Gierlinger N (2018b) Multivariate unmixing approaches on Raman images of plant cell walls: New insights or overinterpretation of results? Plant Methods 14: 52

Prinsloo LC, Du Plooy W, Van Der Merwe C (2004) Raman spectroscopic study of the epicuticular wax layer of mature mango (Mangifera indica) fruit. J Raman Spectrosc 35: 561–567

Renault H, Alber A, Horst NA, Basilio Lopes A, Fich EA, Kriegshauser L, Wiedemann G, Ullmann P, Herrgott L, Erhardt M, et al (2017) A phenol-enriched cuticle is ancestral to lignin evolution in land plants. Nat Commun 8: 14713

Reynoud N, Petit J, Bres C, Lahaye M, Rothan C, Marion D, Bakan B (2021) The Complex Architecture of Plant Cuticles and Its Relation to Multiple Biological Functions. Front Plant Sci 12: 2907

Richter S, Müssig J, Gierlinger N (2011) Functional plant cell wall design revealed by the Raman imaging approach. Planta 233: 763–772

Sasani N, Bock P, Felhofer M, Gierlinger N (2021) Raman imaging reveals in-situ microchemistry of cuticle and epidermis of spruce needles. Plant Methods 17: 1–15

Schmidt U, Hild S, Ibach W, Hollricher O (2005) Characterization of thin polymer films on the nanometer scale with confocal Raman AFM. Macromol Symp 230: 133–143

Schreiber L, Schönherr J (1990) Phase transitions and thermal expansion coefficients of plant cuticles - The effects of temperature on structure and function. Planta 182: 186–193

Segado P, Domínguez E, Heredia A (2016) Ultrastructure of the epidermal cell wall and cuticle of tomato fruit (Solanum lycopersicum L.) during development. Plant Physiol 170: 935–946

Segado P, Heredia-Guerrero JA, Heredia A, Domínguez E (2020) Cutinsomes and CUTIN SYNTHASE1 function sequentially in tomato fruit cutin deposition. Plant Physiol 183: 1622–1637

van Stokkum IH, Mullen K, Mihaleva V (2009) Global analysis of multiple gas chromatography-mass spectrometry (GC/MS) data sets: A method for resolution of co-eluting components with comparison to MCR-ALS. Chemom Intell Lab Syst 95: 150–163

Synytsya A, Čopíková J, Matějka P, Machovič V (2003) Fourier transform Raman and infrared spectroscopy of pectins. Carbohydr Polym 54: 97–106

Szymańska-Chargot M, Pieczywek PM, Chylińska M, Zdunek A (2016) Hyperspectral image analysis of Raman maps of plant cell walls for blind spectra characterization by nonnegative matrix factorization algorithm. Chemom Intell Lab Syst 151: 136–145

Vogg G, Fischer S, Leide J, Emmanuel E, Jetter R, Levy AA, Riederer M (2004) Tomato fruit cuticular waxes and their effects on transpiration barrier properties: Functional characterization of a mutant deficient in a very-long-chain fatty acid β-ketoacyl-CoA synthase. J Exp Bot 55: 1401–1410

Wiley JH, Atalla RH (1987) Band assignments in the raman spectra of celluloses. Carbohydr Res 160: 113–129

Yu MML, Konorov SO, Schulze HG, Blades MW, Turner RFB, Jetter R (2008) In situ analysis by microspectroscopy reveals triterpenoid compositional patterns within leaf cuticles of Prunus laurocerasus. Planta 227: 823–834

Yu MML, Schulze HG, Jetter R, Blades MW, Turner RFB (2007) Raman microspectroscopic analysis of triterpenoids found in plant cuticles. Appl Spectrosc 61: 32–37

